# An inhalation anaesthesia approach for neonatal mice allowing streamlined stereotactic injection in the brain

**DOI:** 10.1101/2020.05.06.078584

**Authors:** Hinze Ho, Adam Fowle, Marisa Coetzee, Ingo H. Greger, Jake F. Watson

## Abstract

Investigating brain function requires tools and techniques to visualise, modify and manipulate neuronal tissue. One powerful and popular method is intracerebral injection of customised viruses, allowing expression of exogenous transgenes. This technique is a standard procedure for adult mice, and is used by laboratories worldwide. Use of neonatal animals in scientific research allows investigation of developing tissues, and enables long-term study of cell populations. However, procedures on neonatal mice are more challenging, due to the lack of reliable methods and apparatus for anaesthesia of these animals. Here, we report an inhalation-based protocol for anaesthesia of neonatal (P0-2) mice, and present a custom 3D-printed apparatus for maintenance of anaesthesia during surgical procedures. This approach significantly enhances animal welfare and facilitates wider and simpler use of neonatal rodents in scientific research. Our optimised method of anaesthesia enables a rapid method of stereotactic injection in neonatal mice for transduction of brain tissue. We demonstrate this procedure for targeted labelling of specific brain regions, and *in vivo* modification of tissue prior to organotypic culture. This anaesthetic approach can be readily employed by any laboratory, and will enable safer use of neonatal rodents across a diverse spectrum of scientific disciplines.

**Highlights:**

- Development of inhalation-based anaesthesia for early postnatal (P0-2) mice
- 3D-printed mould allows anaesthetic maintenance for neonatal surgery
- Improved mouse welfare through reliable neonatal inhalation anaesthesia
- Rapid procedure for brain transduction of mouse litter in under 2 hours

## 1. Introduction

Understanding the mechanisms of brain function requires approaches that reveal the structure and activity of neuronal networks. Recombinant DNA technology has enabled development of powerful genetically encoded tools for visualisation of cell populations, editing of genomic DNA, and tracing of neuronal circuits. These tools can be delivered in repurposed viruses into specific brain areas of adult rodents using intracerebral stereotactic injections (Cetin et al., 2006; Sun and Schaffer, 2018). This widely performed technique involves surgically opening the skull before injection, requiring approximately 40 mins per animal (Cetin et al., 2006), but is highly efficient and provides reliable neuronal transduction in adult animals.

Research using neonatal rodents is less prevalent, but no less valuable, allowing analysis of developmental processes, long-term transgene expression and use of young tissue for sensitive electrophysiology experiments. However, the use of neonates is complicated by a lack of experimental equipment for animals of this size, and critically, the difficulty of anaesthesia in these animals. Neonatal anaesthesia is notoriously unreliable and uncontrollable, posing a significant welfare challenge to their use. Historically, anaesthesia has been performed using hypothermia, where pups are incubated on ice for around 15 minutes, but the depth of hypothermic anaesthesia is unknown and difficult to control. It is unclear whether hypothermia produces anaesthesia or simply immobilisation (Flecknell, 2009), and has been suggested that that anaesthetic recovery is painful and damaging. Therefore, in accordance with 3Rs principles, this approach is now being restricted (Herrmann and Flecknell, 2019).

Here, we present a procedure for inhalation anaesthesia of neonatal mice at P0-2 (**Figure 1A**). Using a 3D-printed apparatus for anaesthetic delivery, we detail a protocol for the controlled maintenance and reliable recovery from inhalation anaesthesia using isoflurane. This method can be readily applied by any laboratory and provides important improvements in neonatal rodent welfare, facilitating their use in a range of scientific disciplines. Using this approach, we perform intracerebral injection of adeno-associated viruses (AAV) into discrete brain areas of P0-2 mice. A complete litter of postnatal animals can be safely and reliably injected in less than 2 hours, offering a rapid and efficient means for brain region specific transduction.

**Figure 1:**
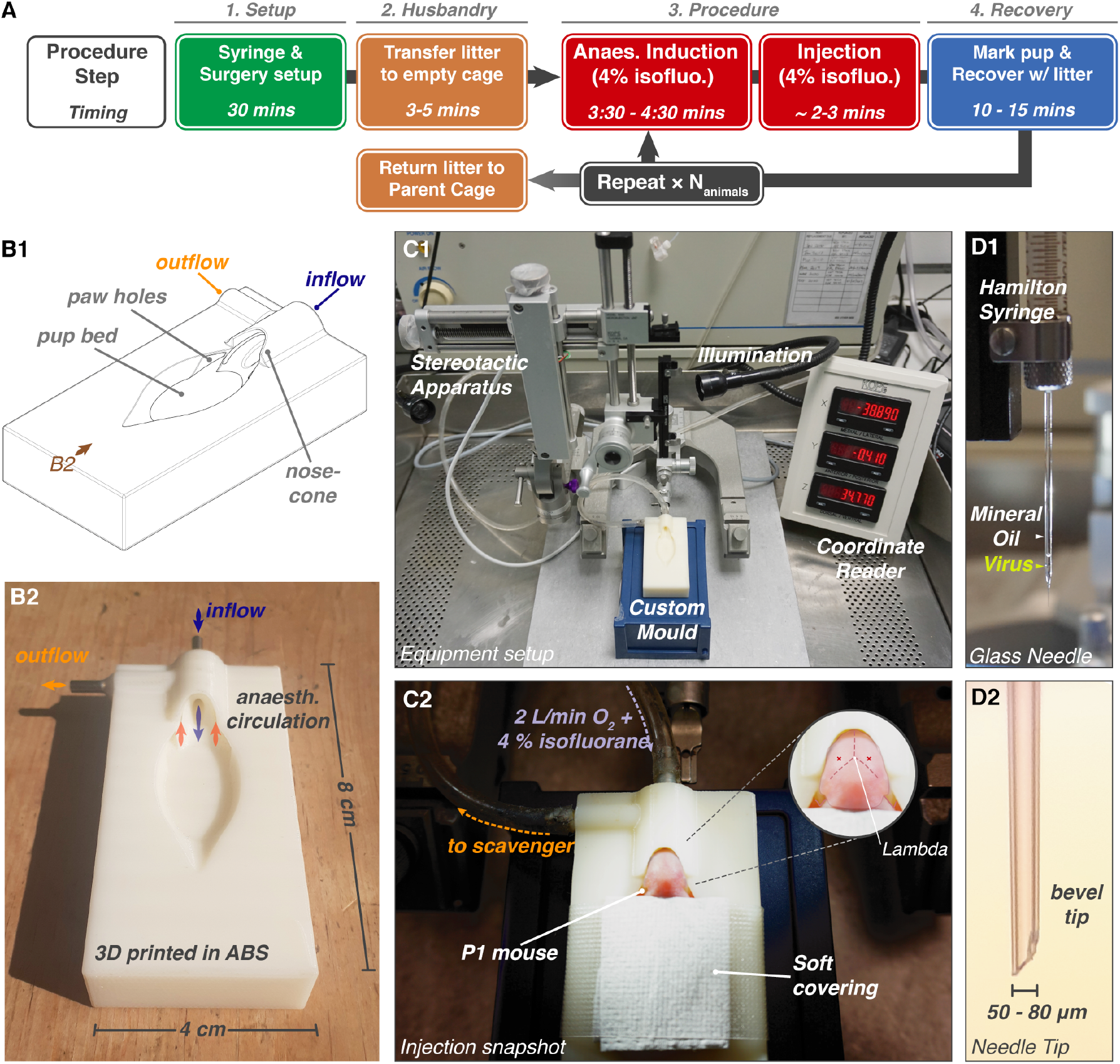
Surgery Procedure and Setup. **A** Procedure flowchart indicating approximate timings. Multiple animals from a litter are injected sequentially, with overlapping recovery periods. **B1** Schematic of anaesthetisation mould. The nose-cone has concentric rings for in (blue) and outflow (orange) of anaesthetic gas. **B2** Mould printed in ABS plastic (view marked in B1). **C1** Procedure setup: a modified adult stereotactic surgery arrangement using the custom neonate mould. Desired height is achieved using an upturned pipette box **C2** P1 mouse during maintenance anaesthesia, covered using a taped medical wipe. Inset: approximate hippocampal injection site (red cross). **D1** Injection uses a glass needle containing mineral oil and minimal viral solution. **D2** Glass needle tip is broken to a 50-80 μm bevelled tip.

## 2. Materials and Methods

### 2.1 Materials

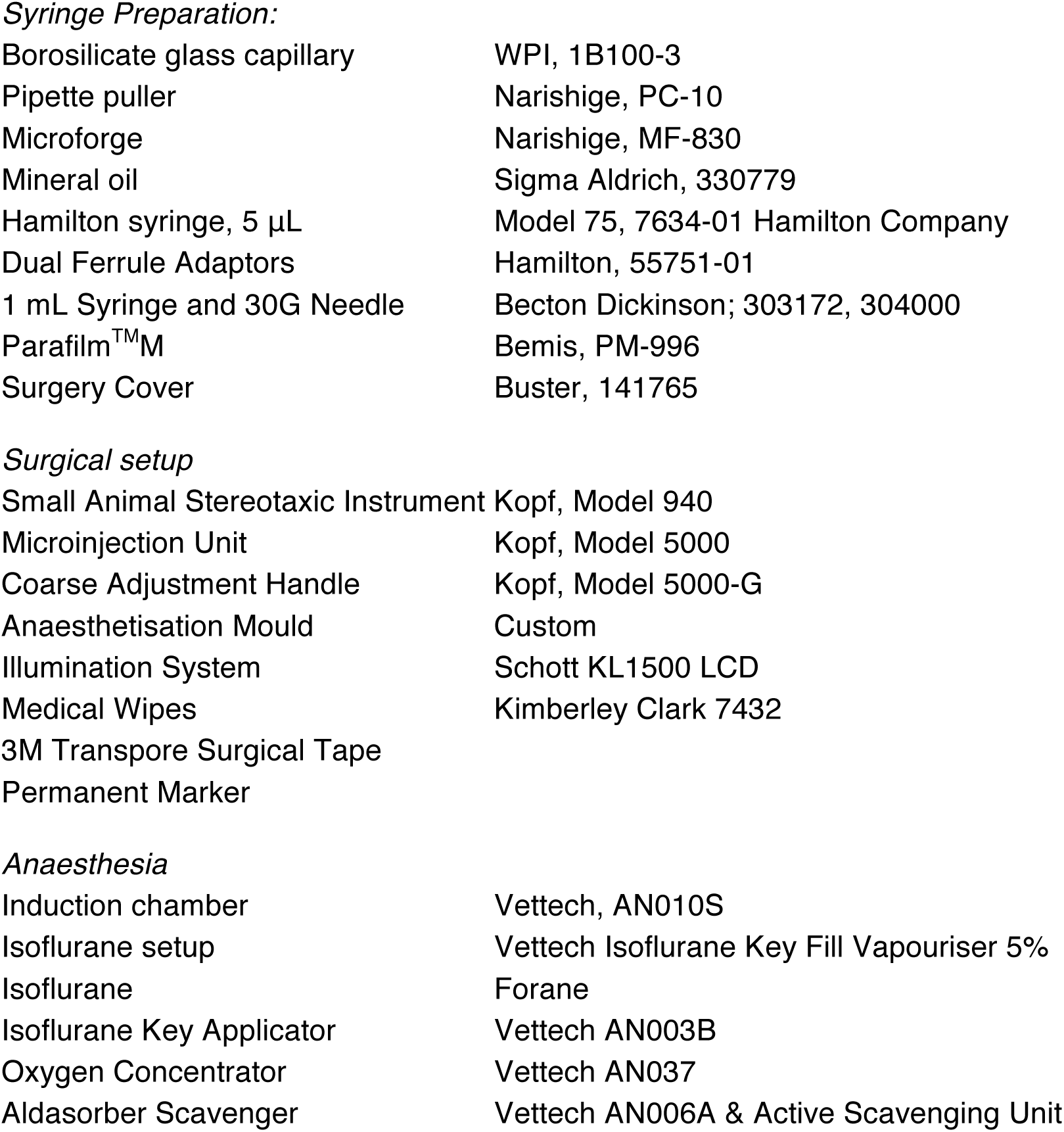

*Syringe Preparation:*
|  |  |
| --- | --- |
| Borosilicate glass capillary | WPI, 1B100-3 |
| Pipette puller | Narishige, PC-10 |
| Microforge | Narishige, MF-830 |
| Mineral oil | Sigma Aldrich, 330779 |
| Hamilton syringe, 5 μL | Model 75, 7634-01 Hamilton Company |
| Dual Ferrule Adaptors | Hamilton, 55751-01 |
| 1 mL Syringe and 30G Needle | Becton Dickinson; 303172, 304000 |
| Parafilm<sup>TM</sup>M | Bemis, PM-996 |
| Surgery Cover | Buster, 141765 |
| *Surgical setup* |  |
| Small Animal Stereotaxic Instrument | Kopf, Model 940 |
| Microinjection Unit | Kopf, Model 5000 |
| Coarse Adjustment Handle | Kopf, Model 5000-G |
| Anaesthetisation Mould | Custom |
| Illumination System | Schott KL1500 LCD |
| Medical Wipes | Kimberley Clark 7432 |
| 3M Transpore Surgical Tape |  |
| Permanent Marker |  |
| *Anaesthesia* |  |
| Induction chamber | Vettech, AN010S |
| Isoflurane setup | Vettech Isoflurane Key Fill Vapouriser 5% |
| Isoflurane | Forane |
| Isoflurane Key Applicator | Vettech AN003B |
| Oxygen Concentrator | Vettech AN037 |
| Aldasorber Scavenger | Vettech AN006A & Active Scavenging Unit |

### 2.2 Animal use

Procedures were performed in accordance with UK Home Office regulations. Experiments conducted in the UK are licensed under the UK Animals (Scientific Procedures) Act of 1986 following AWERB ethical approval. P0-2 wild-type or mutant C57/Bl6 mice were used throughout, and the Gria1-3^fl/fl^ strain was reported previously (Watson et al., 2017). The sex of animals is unascertained.

### 2.3 Viruses

pAAV.CAG.LSL.tdTomato (Addgene #100048) was produced by Addgene, and AAV9.hSyn.HI.eGFP-Cre.WPRE.SV40 (Addgene #105540) was produced by U-Penn Vector Core.

### 2.4 Imaging labelled tissue

Mice were sacrificed at P7-8 and brains were fixed overnight in 4 % paraformaldehyde containing phosphate-buffered saline (PBS) at 4 °C. Tissue was sectioned using a Lecia VT1200 Vibratome, before imaging in PBS using a commercial Leica SP8 confocal microscope.

### 2.5 Organotypic slice preparation

Hippocampi were isolated at P6-8 in high-sucrose Gey’s balanced salt solution (containing in mM: 175 Sucrose, 50 NaCl, 2.5 KCl, 0.85 NaH_2_PO_4_, 0.66 KH_2_PO_4_, 2.7 NaHCO_3_, 0.28 MgSO_4_, 2 MgCl_2_, 0.5 CaCl_2_ and 25 glucose at pH 7.3), cut into 300 μm thick slices using a McIlwain tissue chopper and cultured on Millicell culture inserts (Millipore Ltd) in equilibrated slice culture medium (37 °C/5% CO_2_) containing 78.5 % Minimum Essential Medium (MEM), 15 % heat-inactivated horse serum, 2 % B27 supplement, 25 mM HEPES, 3 mM GlutaMax supplement, 0.25 mM ascorbic acid, with additional 1 mM CaCl_2_ and 1 mM MgSO_4_ (all Thermo Fisher Scientific) (see Watson et al., 2017).

### 2.6 Immunofluorescence

Slices were fixed overnight in PBS containing 4 % paraformaldehyde at 4 °C, before embedding in 2 % agarose for resectioning to 40 μm thickness using a Leica Vibratome (Lee et al., 2015). Sections were permeabilised with 0.5 % Triton X-100 (Sigma-Aldrich) for 2 hours, treated with 1 mg/ml pepsin in 0.2 N HCl for 3 mins, blocked for 3 hours in PBS containing 10 % normal goat serum (NGS) (Sigma-Aldrich), before incubation with anti-GluA2 (RRID:AB_10761960, Sigma Aldrich, 1:500 dilution) and anti-TARP γ8 (RRID:AB_2572272, Frontiers Institute, 1:200) antibodies in 10 % NGS-PBS overnight at 4 °C. After washing in PBS for 2 hours, secondary antibody labelling with anti-Guinea Pig AF647 (RRID:AB_2735091, Invitrogen, 1:500) and anti-Rabbit CF568 (RRID:AB_2833002, Sigma-Aldrich, 1:500) was performed overnight at 4 °C in 10 % NGS-PBS. Washing was repeated before mounting between glass coverslips in ProLong Diamond Mountant (Invitrogen), and imaging using a Leica SP8 confocal microscope.

### 2.7 Electrophysiology

Organotypic cultures at 7-10 days *in vitro* (DIV) were submerged in aCSF containing (in mM): 125 NaCl, 2.5 KCl, 1.25 NaH_2_PO_4_, 25 NaHCO_3_, 10 glucose, 1 sodium pyruvate, 4 CaCl_2_, 4 MgCl_2_, 0.01 SR-95531 and 0.002 2-chloroadenosine (pH 7.3), saturated with 95 % O_2_/5 % CO_2_ (drugs from Tocris Bioscience). 3–6 MΩ borosilicate pipettes were filled with intracellular solution (containing in mM: 135 CH_3_SO_3_H, 135 CsOH, 4 NaCl, 2 MgCl_2_, 10 HEPES, 4 Na_2_-ATP, 0.4 Na-GTP, 0.15 spermine, 0.6 EGTA, 0.1 CaCl_2_, at pH 7.25) for dual voltage-clamp recording (−60 mV holding potential) of neighbouring EGFP positive and negative cells. Schaffer collateral excitatory postsynaptic currents (EPSCs) were evoked using a monopolar glass electrode in CA1 stratum radiatum. Recordings were collected using a Multiclamp 700B amplifier and digitised using a Digidata 1440A interface (Axon Instruments).

### 2.8 Custom mould production

Design was performed using Solidworks®, before conversion to STL file format and printing in acrylonitrile butadiene styrene (ABS) on an Objet Dimension 3D printer. In/outflow pipes made of stainless steel hypodermic tubing were set in drilled holes using Araldite adhesive (**Figure 1B**).

## 3. Procedure for Stereotactic Injection

### 3.1 Setup & preparation

Injection is performed using a Small Animal Stereotactic Frame, with the Microinjection Adaptor for viral delivery using a Hamilton syringe (**Figure 1 C1**). A coarse adjustment handle is recommended for ease of viral delivery. The anaesthetisation mould is positioned below the frame, with the desired height achieved using an upturned pipette box. The inflow pipe is connected to the isoflurane delivery system, and the outflow to a scavenging unit using flexible tubing, and sealed with Parafilm (**Figure 1 C2**).

### 3.2 Syringe Preparation

Borosilicate glass capillaries are pulled into tapered glass needles using a single-step heating protocol on a pipette puller (**Figure 1D**). Bevelled tips are created by breaking the glass using forceps at a diameter of ~50 to 80 μm (**Figure 1 D2**). Tip dimensions of every pipette are confirmed under magnification (e.g. using a microforge).

Excluding air from the syringe ensures accurate volume dispensing; therefore the syringe is entirely filled with mineral oil. The needle is backfilled using a Luer syringe and 30G needle, secured in the Hamilton syringe using glass pipette adaptors, and positioned on the frame.

AAV is loaded into the glass needle through the tip to minimise the wasted viral volume. A drop of AAV-containing PBS is placed on Parafilm beneath the tip and the Hamilton syringe plunger is retracted using the stereotactic frame to draw the required volume into the needle (**Figure 1 D1**). The AAV volume should not exceed that of the glass pipette to prevent cross-contamination between experiments. Glass needles are replaced after injecting 3 animals to prevent clogging or bluntening.

### 3.3 Induction of Anaesthesia

The entire nest, including pups and nesting material is moved from the parent cage to an empty cage containing a surgical liner immediately prior to starting the procedure. Induction of anaesthesia is conducted in a small induction chamber with 2 L/min flow of 4 % isoflurane in O_2_ for 3 to 4.5 mins. After 3 mins, anaesthesia is tested at 30-second intervals by performing a gentle foot pinch using forceps. Loss of motor response denotes sufficient induction. The neonate is then transferred to the anesthetisation mould with inflow of 4 % isoflurane at 2 L/min, secured in place using a tissue covering and micropore tape (**Figure 1 C1**). To minimise exposure of the experimenter to isoflurane and animal allergens, performing the procedure on a downdraft table is recommended.

### 3.4 Intracerebral Injection

Injection positions for specific brain regions are measured relative to lambda (**Figure 1 C2** insert). Under LED illumination, the pipette is positioned above the injection site, and depth axis (z) is zeroed at the height of the skin. Injection is performed by rapidly and firmly lowering the glass needle to penetrate both the skin and skull. After penetration, the injection depth is adjusted to the desired location, and virus is slowly dispensed under hand control. We use 0.5 μL injection of virus at around 1×10^12^ GC ml^−1^ per hemisphere. The glass pipette is retracted gently from the skull, and injection is repeated on the second hemisphere if required.

### 3.5 Recovery

The total duration of anaesthesia is approximately 6-10 mins. After injection, the animal’s tail is marked with pen to discriminate from uninjected animals, and returned to the nest with the remainder of the litter for recovery from anaesthesia. Subsequent animals are injected following the same procedure. Once all injections have been completed, pups are checked for successful recovery (usually 10-20 minutes), as determined by spontaneous limb mobility. The nest is returned to the parent cage after successful recovery of all animals. Injection of a litter of 6-8 pups usually takes around 1.5 to 2 hours from setup to recovery.

## 4. Results

### 4.1 Induction of neonatal inhalation anaesthesia

We aimed to develop a robust and reliable technique for anaesthetising neonatal mice, as an alternative to hypothermia. We monitored the induction of anaesthesia of P0-2 mice using 4% isoflurane in an induction chamber, as is typically used for anaesthesia of adult mice. At least 3 minutes of isoflurane exposure was required for reliable and sufficient anaesthetic depth, which is significantly longer than for adult mice. However, the duration must be confirmed on every animal using a gentle foot pinch, as movement levels of neonatal mice can be minimal even without complete anaesthetic induction.

We have noticed that increased duration of anaesthesia can correlate with non-recovery, however this has not been experimentally examined for ethical reasons. Due to this observation, we induce pups for no more than 4′30″, and if response to foot pinch is still evident, the animal will not undergo the procedure. Using induction of between 3′ to 4′30″, a trained user can expect minimal non-recovery occurrences (30/30 mice recovered across 7 injected litters).

### 4.2 Maintenance of anaesthesia using a custom frame

Maintenance of anaesthesia requires apparatus for circulation of anaesthetic at the animal’s snout, while allowing access to the cranium. We designed and produced a 3D-printed gas flow apparatus consisting of a neonate-shaped mould with a nose-cone formed of concentric rings for anaesthetic delivery. The inflow (central) and outflow (outer) routes are connected to standard anaesthetic apparatus via metal adaptors that are embedded in the frame (**Figure 1B**). The STL file for production of this apparatus will be freely available for academic use upon request.

### 4.3 Consideration of neonatal mouse age

To limit the loss of injected pups, in particular from maternal cannibalism, we and others (Li and Daly, 2002) perform injection at P1 rather than P0, as infant mortality occurring in the first 24 hours post birth is avoided.

For adult animals, access to the brain requires drilling through the skull, while at P0-2, the soft skull can be penetrated by a glass injection needle. This greatly simplifies the procedure, dramatically decreasing the protocol duration, and allows injection of a complete litter (6-8 pups) in around 1.5 - 2 hours from setup to recovery. Due to the developmental thickening of the skull, this approach can only be used for animals up to P3.

### 4.4 Intracerebral injection for neuronal labelling

We demonstrate this procedure for two experimental approaches. Firstly, we injected two discrete brain areas for sparse fluorescent labelling of neuronal populations by coinjection of AAVs expressing Cre-dependent tdTomato (lox-Stop-lox cassette) and Cre-EGFP at a ratio of 10,000:1 (**Figure 2A**, see Weiler et al., 2018). 0.5 μl of viral mixture was injected per hemisphere at a total concentration of 1.5 ×10^12^ GCml^−1^ into either the hippocampus (From lambda, in mm: Rostral (+)/ caudal (−): 0.0, Lateral: ± 1.3, Ventral: 1.4) or frontal cortex (Rostral (+)/ caudal (−): +1.5, Lateral: ± 1.0, Ventral: 1.0). On brain dissection at P8, robust expression was evident in the corresponding brain areas (**Figure 2B-C**).

**Figure 2:**
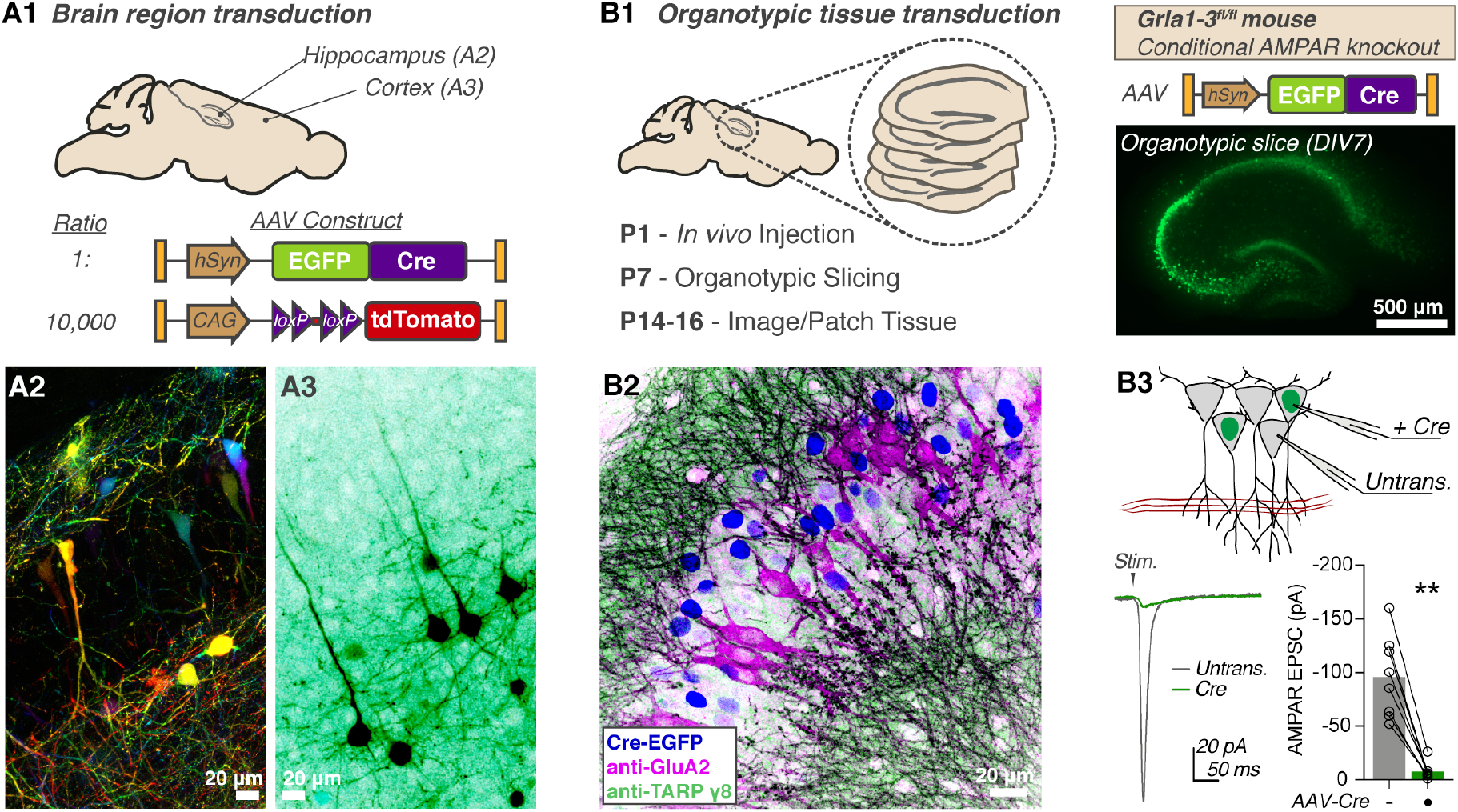
Stereotactic transduction of neonatal brain tissue. **A** Sparse fluorescent labelling is achieved by coinjection of Cre-dependent tdTomato, and Cre-EGFP expressing AAVs (**A1**). Representative images of tdTomato fluorescence from hippocampus (**A2**, coloured by z-depth) and frontal cortex (**A3**, inverted LUT). **B1** P1 injection of Cre-AAV in Gria1-3^fl/fl^ mice allows *in vivo* hippocampal AMPAR knockout before organotypic slice preparation. **B2** Knockout is confirmed by immunofluorescent staining of the CA3 area showing cells mutually exclusive for GluA2 or nuclear-localised Cre (Nuclear Cre-GFP: blue; anti-GluA2: magenta; anti-TARP g8: green), and (**B3**) dual electrophysiological recording of CA1 evoked synaptic currents, which is abolished in organotypic slices 14 days after *in vivo* Cre transduction (Untrans: −95.71 ±13.34 pA; Cre: −7.86 ±2.75 pA; n=8, Wilcoxon matched-pairs signed rank test, p=0.0078). Recording configuration and example trace are depicted.

### 4.5 Intracerebral injection for organotypic tissue preparation

Hippocampal organotypic slice culture is a powerful method allowing *ex vivo* analysis of neuronal physiology (Gähwiler et al., 1997), but requires tissue harvesting at around P7. Neonatal injection allows *in vivo* transduction of tissue prior to harvesting, increasing the flexibility of experimental timings. This is particularly important for genomic manipulations such as Cre-dependent gene excision, which can require weeks for removal of highly abundant or stable target proteins. For instance, the AMPA receptor (AMPAR), a glutamate-gated synaptic receptor, requires at least 14 days from Cre-DNA transduction before complete protein loss is achieved in conditional knockout mice (Lu et al., 2009).

To optimise organotypic study of AMPAR knockout tissue, we performed injection of AAV-Cre-EGFP (0.5 μl at 3 ×10^12^ GCml^−1^) into the hippocampus of P1 Gria1-3^fl/fl^ mice (Conditional *GRIA1-3* knockout). Cre transduction will prevent expression of the three AMPAR subunits present in hippocampal pyramidal cells. When tissue was dissected at P7, robust EGFP expression was seen in organotypic slices (**Figure 2 B1**). These slices were cultured until DIV8 for analysis. At this time-point, immunofluorescent detection of AMPARs (antibody to GluA2 protein) in fixed slices showed a mosaic pattern, with AMPAR expression specifically absent from Cre-expressing CA3 cells (**Figure 2 B2**). Similarly, dual patch-clamp recording of synaptic currents at CA1 pyramidal cells showed an absence of AMPAR currents in Cre-transduced neurons, confirming successful modification of organotypic tissue *in vivo* (**Figure 2 B3**). These examples demonstrate the utility of neonatal injection for modification of developing brain tissue, and add a further tool to the neurophysiologist’s toolkit.

## 5. Discussion

Intracerebral injection of neonatal rodents is a powerful approach for neuroscience research, however reliable anaesthesia of early postnatal animals has been difficult to achieve. Previous reports have been reliant on either hypothermic anaesthesia (Cheetham et al., 2015; Kim et al., 2014), or even a lack of anaesthesia (Li and Daly, 2002), raising major welfare issues (Herrmann and Flecknell, 2019).

Our inhalation-based technique offers a refinement of neonatal rodent anaesthesia that can be employed for recovery procedures on early postnatal animals. Induction of anaesthesia is performed in an induction chamber, however, due to the rapid elimination of isoflurane from the body, maintenance of anaesthesia requires constant delivery throughout the procedure. Apparatus for neonatal stereotactic injection has been reported, but is designed for hypothermic anaesthesia (Cunningham and McKay, 1993). We therefore designed a 3D printable mould for delivery of inhalational anaesthetics. 3D printing allows simple size-scaling of the apparatus for application to various ages or rodent species, and offers affordable production requirements. This anaesthetic procedure can be used not only for intracerebral injections, but any other cranial procedures, and simple redesign of the printed apparatus would allow access to other regions of the animal, for surgical procedures across a range of scientific disciplines. We report the use of isoflurane, which has no negative effects on cognition from brief exposure (Rosenholm et al., 2017), however the procedure can be applied to other inhalational anaesthetics.

We have demonstrated the value of neonatal intracerebral injection for *in vivo* tissue transduction before organotypic culture, and have previously employed this approach in synaptic receptor research (Watson et al., 2017). Our technique offers a rapid and efficient protocol for neuronal transduction, and published neonatal brain maps (Pilpel et al., 2009) will aid stereotactic targeting of any brain area. The protocol and apparatus that we report presents a significant improvement in rodent welfare, and can be easily adapted to facilitate greater use of neonatal animals in both neuroscience and wider scientific research.

## Acknowledgements

The authors are deeply grateful to the Biological Services team at both the Laboratory of Molecular Biology and Ares facilities for supporting this work, in particular Lesley Drynan and Richard Berks who facilitated its setup. We would also like to thank Ana González Rueda for initial assistance with neonatal injections, Txomin Lalanne for consultation on initial mould design, and Steve Scotcher for assistance with 3D printing. We are very grateful to Alexandra Pinggera and Julia Morud Lekholm for critical reading of the manuscript and constructive feedback. This work was supported by the Medical Research Council (MC_U105174197).

